# Critical information thresholds underlying concurrent face recognition functions

**DOI:** 10.1101/2020.06.22.163584

**Authors:** Genevieve L. Quek, Bruno Rossion, Joan Liu-Shuang

## Abstract

Humans rapidly and automatically recognise faces on multiple different levels, yet little is known about how the brain achieves these manifold categorisations concurrently. We bring a new perspective to this emerging issue by probing the relative informational dependencies of two of the most important aspects of human face processing: categorisation of the stimulus *as a face* (generic face recognition) and categorisation of its familiarity (familiar face recognition). Recording electrophysiological responses to a large set of natural images progressively increasing in image duration (Expt. 1) or spatial frequency content (Expt. 2), we contrasted critical sensory thresholds for these recognition functions as driven by the same face encounters. Across both manipulations, individual observer thresholds were consistently lower for distinguishing faces from other objects than for distinguishing familiar from unfamiliar faces. Moreover, familiar face recognition displayed marked inter-individual variability compared to generic face recognition, with no systematic relationship evident between the two thresholds. Scalp activation was also more strongly right-lateralised at the generic face recognition threshold than at the familiar face recognition threshold. These results suggest that high-level recognition of a face *as a face* arises based on minimal sensory input (i.e., very brief exposures/coarse resolutions), predominantly in right hemisphere regions. In contrast, the amount of additional sensory evidence required to access face familiarity is highly idiosyncratic and recruits wider neural networks. These findings underscore the neurofunctional distinctions between these two recognition functions, and constitute an important step forward in understanding how the human brain recognises various dimensions of a face in parallel.

**Significance Statement:** The relational dynamics between different aspects of face recognition are not yet well understood. We report relative informational dependencies for two concurrent, ecologically relevant face recognition functions: distinguishing faces from objects, and recognising people we know. Our electrophysiological data show that for a given face encounter, the human brain requires less sensory input to categorise that stimulus as a face than to recognise whether the face is familiar. Moreover, where sensory thresholds for distinguishing faces from objects are remarkably consistent across observers, they vary widely for familiar face recognition. These findings shed new light on the multifaceted nature of human face recognition by painting a more comprehensive picture of the concurrent evidence accumulation processes initiated by seeing a face.

## Main Text

Faces hold an exceptional status in the human brain, conveying a great deal of meaningful social information that is recognised nearly effortlessly by neurotypical adults. Yet the relative ease of face recognition belies the complex and multifaceted nature of this key human faculty, which comprises a heterogeneous set of processes that culminate in functional categorisations of a face *as a face*, its sex, emotion, familiarity, identity (via associated semantic attributes/names), and beyond (1). Remarkably, these various functions appear to be both concurrent and outside volitional control, such that to encounter a face is to almost instantaneously ‘recognise’ it in a multitude of different ways.

Despite the concurrent and multifaceted nature of face recognition/categorization^*^, there is a long tradition of studying its various facets in isolation, by having observers view face images that differ along a single dimension (e.g., familiarity status) and inspecting how that manipulation modulates a behavioural or neural response (2–5). Within this modular framework, empirically relating the different levels of face categorisation to one another necessitates contrasting responses across observer tasks, and therefore different face encounters (e.g., response times (RTs) for recognising faces amongst objects *vs.* RTs for recognising familiar faces amid unfamiliar ones) (6–11). Since each task is associated with specific goals, distractor images, face stimuli, and/or observer strategies, general experimental confounds hamper the identification of distinctive characteristics associated with the various face categorisation functions. As such, this cross-task comparative approach offers limited insight into the relational dynamics between the various categorisation processes that are elicited in parallel by the same face encounter.

Recently, however, a new wave of face research has emerged aimed at elucidating how the brain extracts information along different face dimensions *concurrently* (12–15). In contrast to second-order comparisons of face functions (i.e., contrasts across different tasks/face encounters), this approach investigates the different levels of categorisation reflected in the exact same neural response elicited by a given face encounter, often by applying multivariate pattern analysis (MVPA) techniques to high temporal resolution electro/magneto-encephalographic data (EEG, MEG) (16). For example, a recent MEG study used this approach to examine the temporal dynamics underlying concurrent categorisation of face images along the dimensions of familiarity, sex, and age. Contrasting the time course of decoding associated with each dimension in the neural response showed that the age and sex of a face were categorised earlier than its identity (14). However, where this multivariate approach focuses near-exclusively on contrasting the relative onset and duration of different categorisations that follow a face presentation, cognitive processes can differ not only in their temporal unfolding, but also in the amount of sensory evidence they require to proceed (17). To date, the latter possibility has received little explicit exploration in the context of multifaceted face categorisation, such that the relative informational dependency of concurrent face categorisation functions remains largely unknown. Yet this aspect is no less important, as attested by the many (modular) investigations of evidence accumulation for the different types of face categorisation that have manipulated image duration (18–21), spatial resolution (22, 23), or visibility (24).

The current study characterised how increasing access to sensory face input influences two concurrent ecologically relevant face functions: *generic face categorisation* (i.e., recognising that a visual stimulus as a face, as opposed to another type of object) and *familiar face categorisation* (i.e., recognising that a face is one you have encountered before). We recorded high-density scalp EEG while observers viewed a large number of widely variable, unsegmented face images in which we constrained sensory evidence by parametrically varying either viewing time (i.e., stimulus presentation duration, Expt. 1, or spatial frequency content (i.e., image resolution, Expt. 2) (22). By measuring the effect of increasing temporal/spatial exposure on implicit neural indices of concurrent generic and familiar face categorisation, we identified the minimal amount of sensory input capable of driving each form of categorisation^†^. Importantly, we inspected these spatiotemporal thresholds at the individual observer level, with the goal of explicitly relating the informational requirements of face categorisation processes within the same observer. On one hand, the two recognition functions may be tightly coupled in the human brain, such that the sensory input diagnostic for recognising a visual stimulus as a face also enables recognition of whether that face is familiar. On this possibility, the distribution of individual observer thresholds for generic face categorisation should vary as a function of face familiarity (i.e., they should be at least partially non-overlapping). On the other hand, if faces can be successfully distinguished from objects based on less sensory input than is required for familiar face recognition, generic face recognition thresholds should be similarly distributed regardless of face familiarity, but reliably offset from familiar face recognition thresholds.

## Results

To quantify the amount of sensory input required for successful generic and familiar face recognition, we used an EEG frequency-tagging paradigm in which observers view rapid sequences of natural object categories (e.g., plants, animals, buildings, vehicles, etc.) with faces embedded at strict periodic intervals (Fig. 1A). Stimulating the visual system in this way yields separable electrophysiological indices of *i)* general visual processing (i.e., processing common to faces and objects, measurable at the frequency of image presentation), and *ii)* category-selective visual processing (i.e., the differential response to faces *vs.* objects, measurable at the frequency of face presentation) that are identifiable at the level of individual observers (22, 25–27). For our purposes here, the category-selective visual response provided an index of generic face recognition, as it can only arise if the neural response evoked by faces in the sequence is both consistently similar across different face exemplars, yet consistently different to the responses evoked by object images. We obtained a corresponding index of familiar face recognition by computing the difference between face-selective responses elicited by sequences containing either highly *Familiar* or *Unfamiliar* faces (Fig. 1A) that varied widely in pose, expression, background, etc. In this way, neither recognition index could be driven by low-level image properties (25, 28). In two separate groups of observers, we tracked both indices as a function of parametric increase in either image presentation duration (Expt. 1, Fig. 1B) or spatial frequency content (Expt. 2; Fig. 1C), in both cases identifying the minimal informational input required for successful generic and familiar face recognition. We defined these thresholds (referred to hereafter as *Gen-*Thresh and *Fam-*Thresh respectively) as the first parametric increment at which the relevant response exceeded a bootstrapped criterion cut-off equivalent to *p* < .01 (one-tailed, see Methods).

**Fig. 1.**
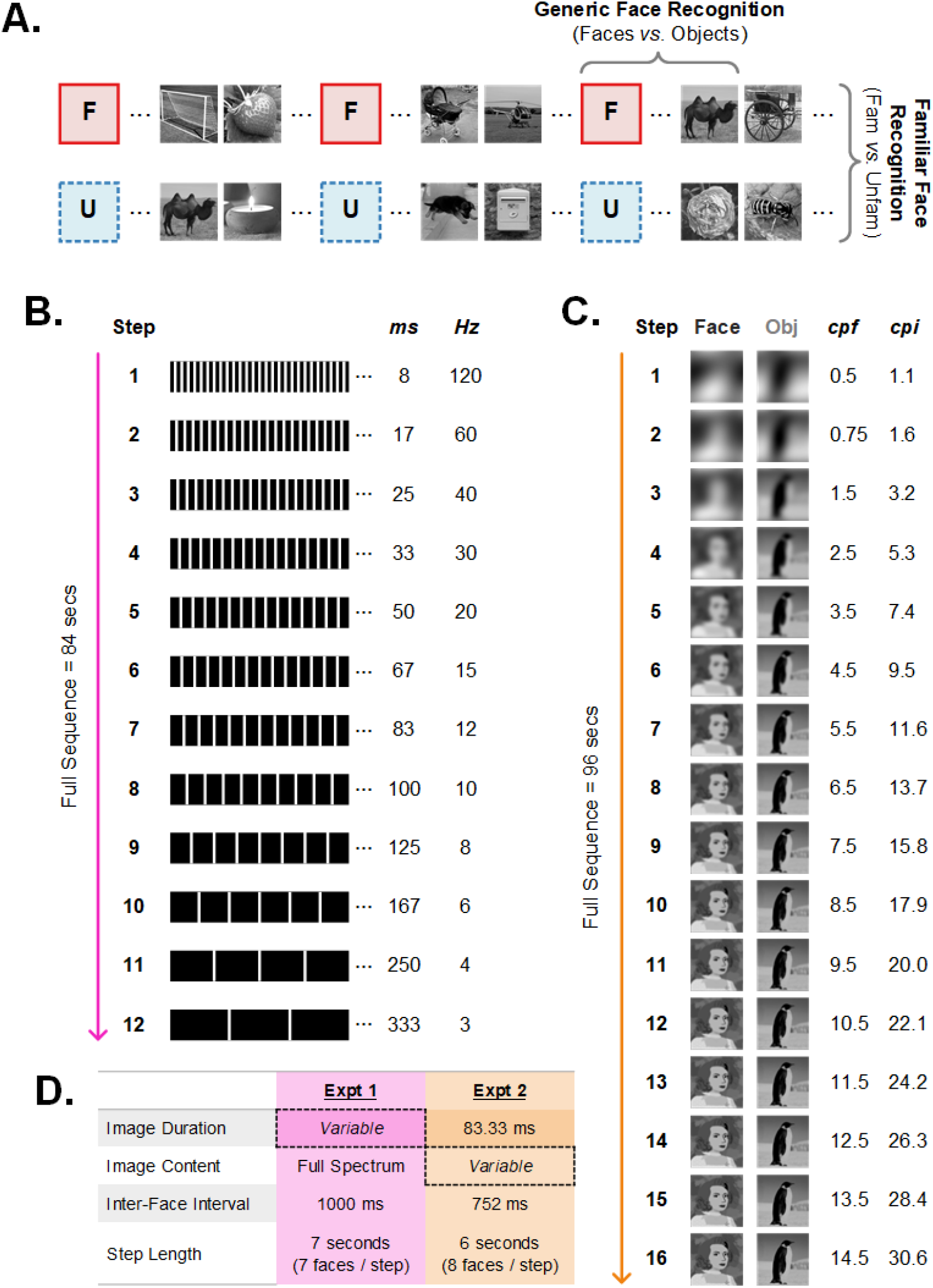
Design overview. **A.** The basic frequency-tagging paradigm consists of a rapid stream of various object categories interspersed with *Familiar* or *Unfamiliar* faces at strictly periodic intervals (indicated here by placeholders F and U) (25). **B**. Sequences in Expt. 1 contained full spectrum images whose duration increased every 7 s. The inter-face interval was always 1000 ms for a total of 84 face presentations/sequence. **C.** In Expt. 2, image duration was fixed at 83.33 ms, with image resolution increasing every 6 s (depicted here with a cartoon face licensed under CC BY-NC-SA 2.0; face images in the real experiment were photographs). The inter-face interval was always 752 ms, for a total of 127 faces/sequence (cpf = cycles per (median) face width; cpi = cycles per image cartoon face shown here **D.** Summary of the differing design properties in Expts. 1 & 2.

### Expt. 1: Increasing Image Duration

In Expt. 1 we examined how generic and familiar face categorisation evolved as a function of increasing image duration over 12 incremental steps (see Methods). At the group-level, inspection of the scalp topographies (Fig. 2A) and response profiles within a predefined bilateral occipitotemporal (OT) ROI revealed that the generic face categorisation responses for *Familiar* and *Unfamiliar* faces emerged gradually, both reaching significance at the same image duration (i.e., 33.33 ms, Fig. 2C). Inspection of the normalised topographical maps indicated stable activation of (right) OT channels throughout all supra-threshold steps (Fig. 2B), validating our *a priori* selection of ROI electrodes (27, 29). In contrast, the familiar face recognition response profile hovered around the noise baseline at the shortest image durations before reaching significance at a slightly longer presentation duration (50 ms) than was observed for generic face recognition (Fig. 2D). Corresponding scalp topographies (bottom row, Fig. 2B) implicated similar bilateral OT regions as for generic face recognition. At the group-level, this familiar face recognition response increased to a local peak at 83 ms image duration, then plateaued until the last duration step, suggesting a sustained differentiation between *Familiar* and *Unfamiliar* faces. However, many individual participant response profiles in fact exhibited a transient discrimination between *Familiar* and *Unfamiliar* faces (Fig. 2E), with little difference between conditions at the longer image durations (e.g., >167ms, see Fig. S1 for all participant profiles).

**Fig. 2.**
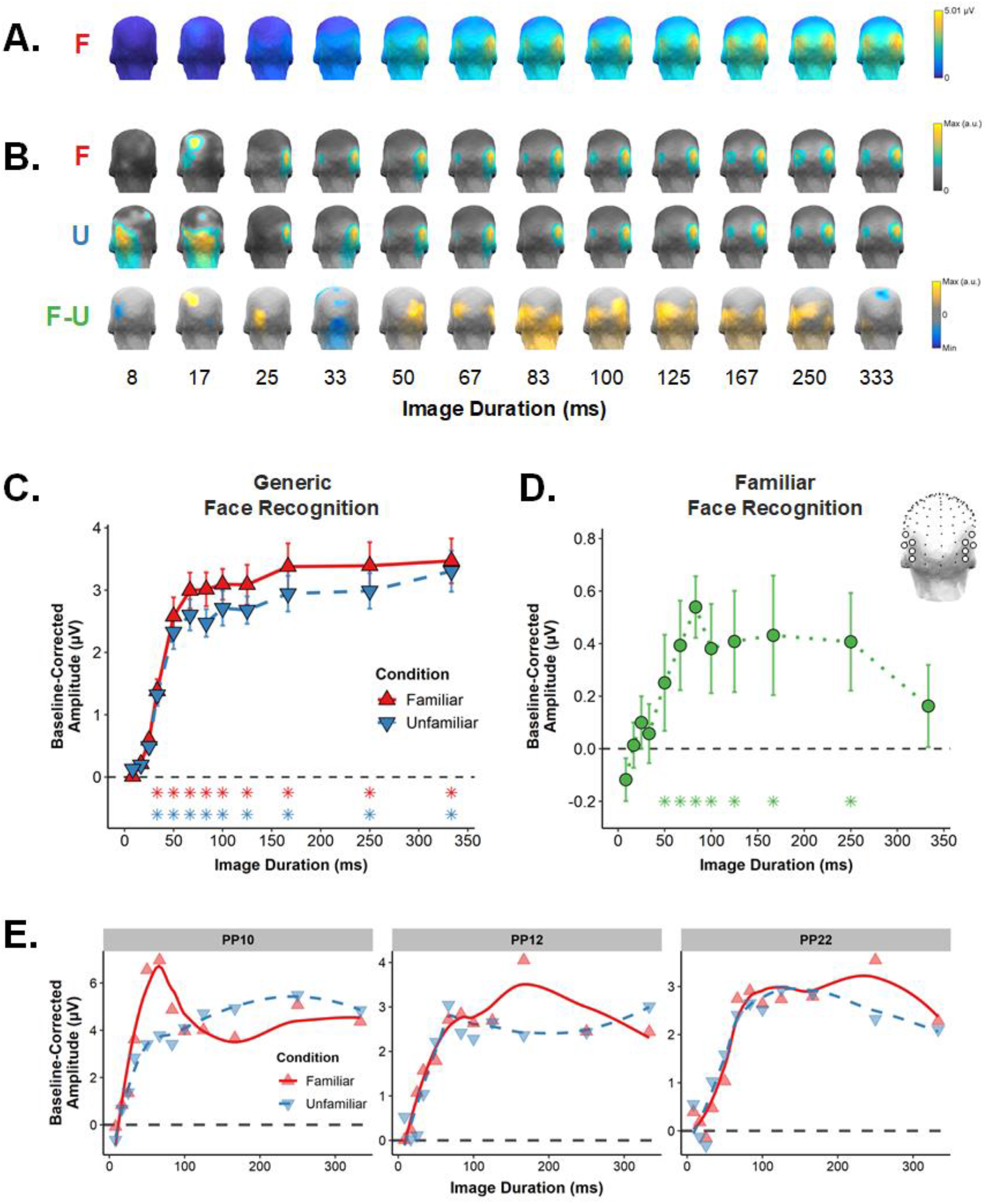
Group-level data for Expt. 1. F = Familiar, U = Unfamiliar. **A.** Baseline-corrected amplitude scalp topographies for the *Familiar* condition show the face-selective response emerging as a function of increasing duration. **B.** Normalised scalp topographies as a function of image duration for all conditions. **C.** The *Familiar* and *Unfamiliar* generic face categorisation response profiles within the OT ROI (right inset). **D.** The familiar face recognition response profile within the same ROI. Error bars are SEM, asterisks correspond to image durations eliciting a significant response (*p* <. 01, one-tailed). **E.** Face recognition response profiles for three example participants, shown with local polynomial regression fits (see Fig. S1 for all individual profiles).

To more precisely characterise the relative informational dependencies of generic and familiar face categorisation, we focused on the distributions of individual-level thresholds for each of these processes (Fig. 3). Generic face categorisation thresholds were distributed very similarly in the *Familiar* and *Unfamiliar* conditions, in both cases being narrow and peaking over 33-50ms image duration. Both Wilcoxon and Kolmogorov-Smirnov (K-S) tests revealed no significant difference between *Gen*-Thresh distributions for *Familiar* and *Unfamiliar* faces (Wilcoxon *p* =.704; K-S *p* = .833). In contrast, familiar face categorisation thresholds were distributed much more broadly, peaking over longer durations of 83-100ms. Nonparametric tests indicated that both central tendency and shape differed significantly between the *Fam-*Thresh and *Gen-*Thresh (averaged across *Familiar* and *Unfamiliar*) distributions (Wilcoxon *p*=.010; K-S *p* = .005), suggesting that familiar face recognition not only necessitates longer temporal exposure than generic face recognition, but that it also depends more on the individual processing efficiency of each observer.

**Fig. 3.**
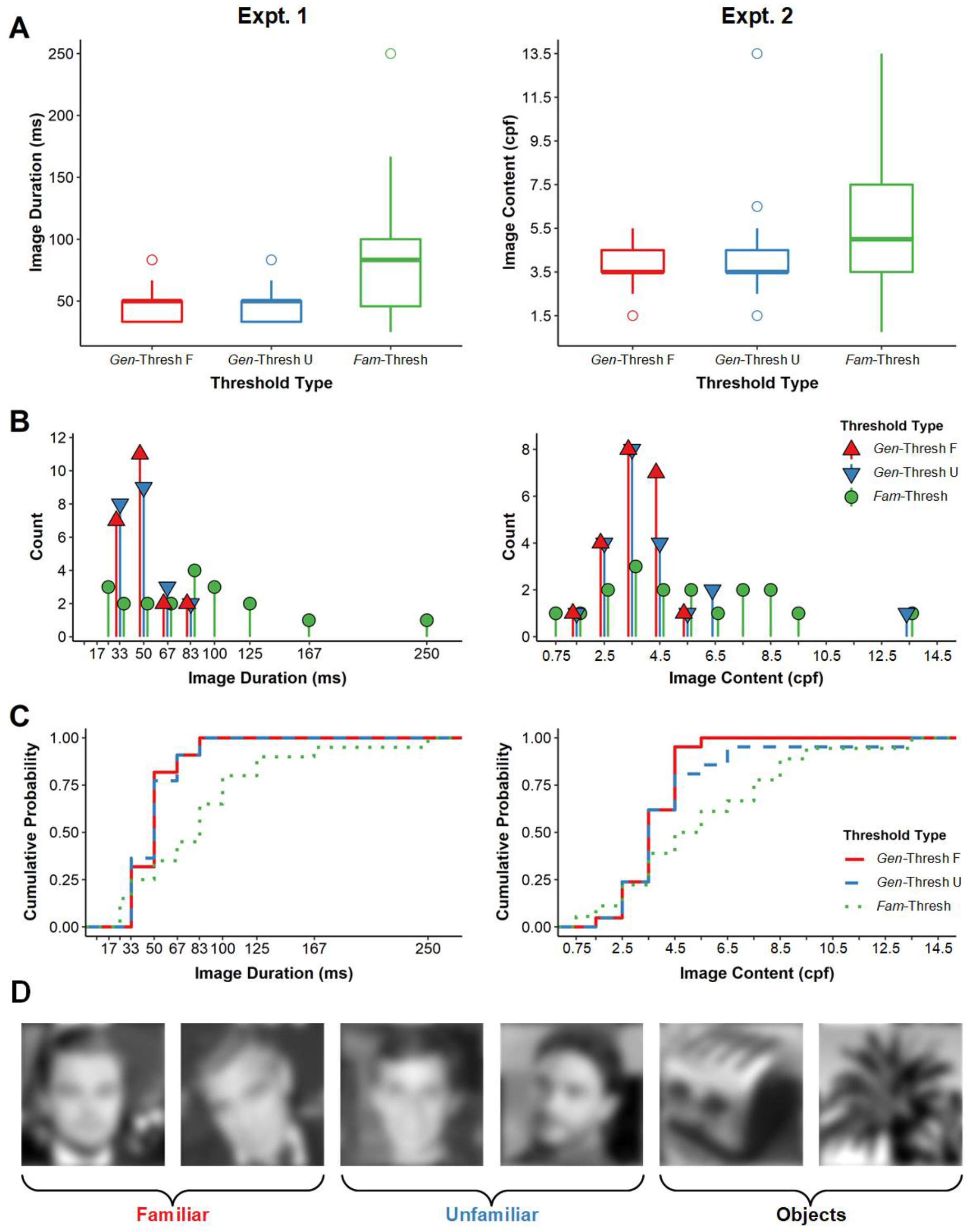
Distribution of individual *Gen*-Thresh and *Fam*-Thresh values in Expt. 1 (left column) and Expt. 2 (right column), represented as **A.** box-and-whisker plots, **B.** Frequency counts, and **C.** Empirical cumulative distribution functions. **D.** Examples of *Familiar*, *Unfamiliar* and Object stimuli at the median *Gen*-Thresh for Expt. 2 (3.5 cpf).

An interesting possibility is that observers may vary reliably terms of their minimal required image duration supporting both generic and familiar face recognition. Somewhat surprisingly, however, we observed no systematic relationship between individual *Gen-*Thresh and *Fam-*Thresh values (*r*_pearson_ = 0.09, *p* = 0.70). As can be seen in Fig. 4A, observers with more efficient generic face recognition did not also tend to exhibit more efficient familiar face recognition. Instead, the two thresholds appeared to vary independently, such that individuals with very similar generic face recognition thresholds displayed markedly different familiar face recognition thresholds.

**Fig. 4.**
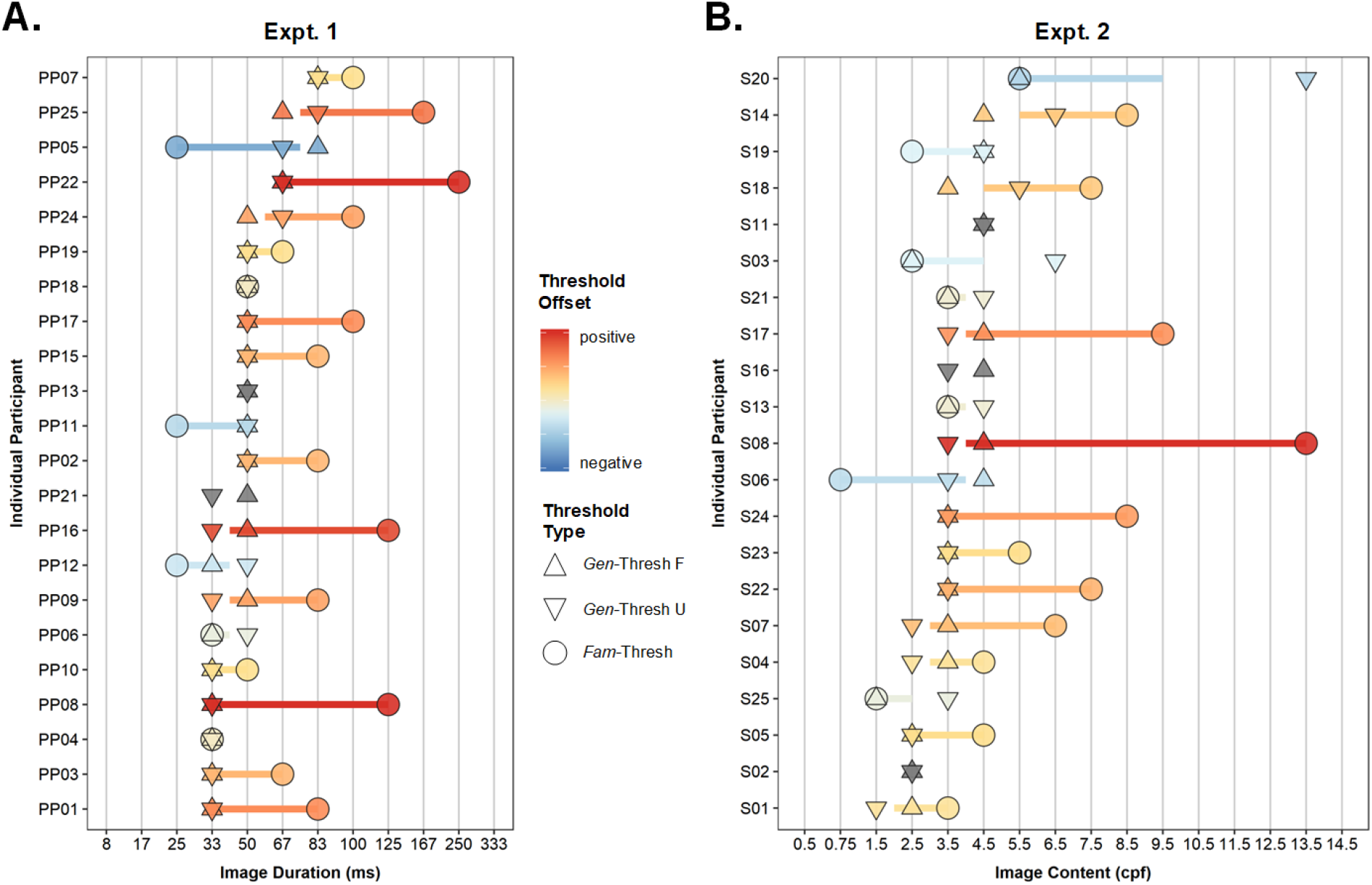
Individual threshold values for Expt. 1 (A) and Expt. 2 (B), rank ordered by average *Gen*-Thresh. Colours indicate the degree and direction of offset between *Gen*-Thresh and *Fam*-Thresh (grey points are observers with no identifiable *Fam*-Thresh).

Interestingly, while the evolving generic and familiar face categorisation responses were both consistently located over OT channels (Fig. 2B), the former appeared to be more strongly right-lateralised, suggesting there may be different underlying neural regions associated with processing critical sensory input required for each recognition function. This pattern was particularly evident in the scalp topographies corresponding to the (individually-defined) threshold points (Fig. 5A), where the lateralisation index (i.e., right ROI – left ROI) was significantly stronger at *Gen-*Thresh than at *Fam*-Thresh, *t*(1,19) = 2.51, *p* < 0.02 Fig. 5B). Further decomposing this pattern revealed that, at the shortest durations supporting the distinction between faces and objects, responses were similarly right-lateralised for both *Familiar* and *Unfamiliar* faces (Fig. 5C). By contrast, at the shortest durations supporting *Familiar/Unfamiliar* face differentiation, *Familiar* faces evoked a more bilateral face-selective response than *Unfamiliar* faces.

**Fig. 5.**
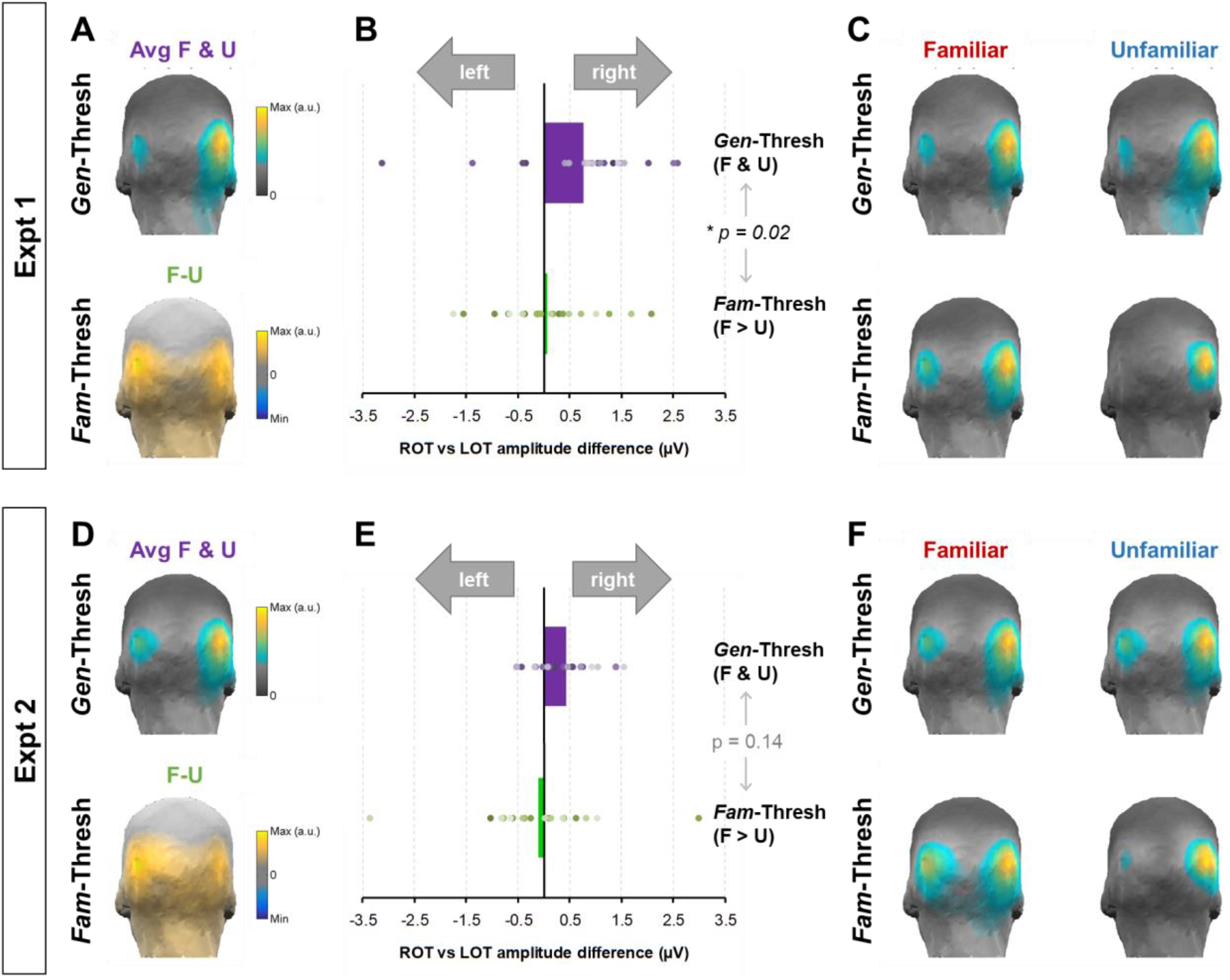
Response lateralisation at *Gen-* and *Fam-*Thresh in Expt.1 (top row) and Expt. 2 (bottom row). **A & D.** Mean normalised topographies at the individual observer values of Fam-Thresh and Gen-Thresh (averaged across Familiar and Unfamiliar). **B & E.** Differences in baseline-corrected amplitudes between the right and left OT ROIs for each threshold type. Bars reflect the group mean; dots are individual observers. **C & F.** Decomposition of the mean normalised topographies at threshold into the *Familiar* and *Unfamiliar* conditions. Scalp activation at *Fam*-Thresh was comparatively more bilateral for *Familiar* faces than *Unfamiliar* ones.

### Expt. 2: Increasing Image Content

In Expt. 2 we examined generic and familiar face recognition responses as a function of increasing spatial frequency (SF) content (see Methods). At the group-level, the face categorisation response emerged at the same coarse image resolution for both *Familiar* and *Unfamiliar* faces (i.e., *Gen*-Thresh = 3.5 cpf or 7.4 cpi) and increased steadily before stabilising around 10.5 cpf (Fig. 6B). As was the case in Expt. 1, inspection of the normalised scalp topographies (Fig. 6A) indicated consistent activation of lateral channels across increasing image content, suggesting that similar OT neural populations were engaged regardless of the spatial frequency content of the face images. Although the group-level face recognition response (Fig. 6C) also rose significantly above noise at 3.5 cpf, this difference between *Familiar* and *Unfamiliar* responses did not appear to stabilise until a slightly higher image resolution (i.e., 5.5 cpf, or 11.6 cpi). Just as in Expt. 1, inspection of the *Familiar* and *Unfamiliar* face responses for individual observers revealed some dissociation with the group-level averaged response profiles, once against bolstering our approach to focus on threshold differences at the individual-level (Fig. S2).

**Fig. 6.**
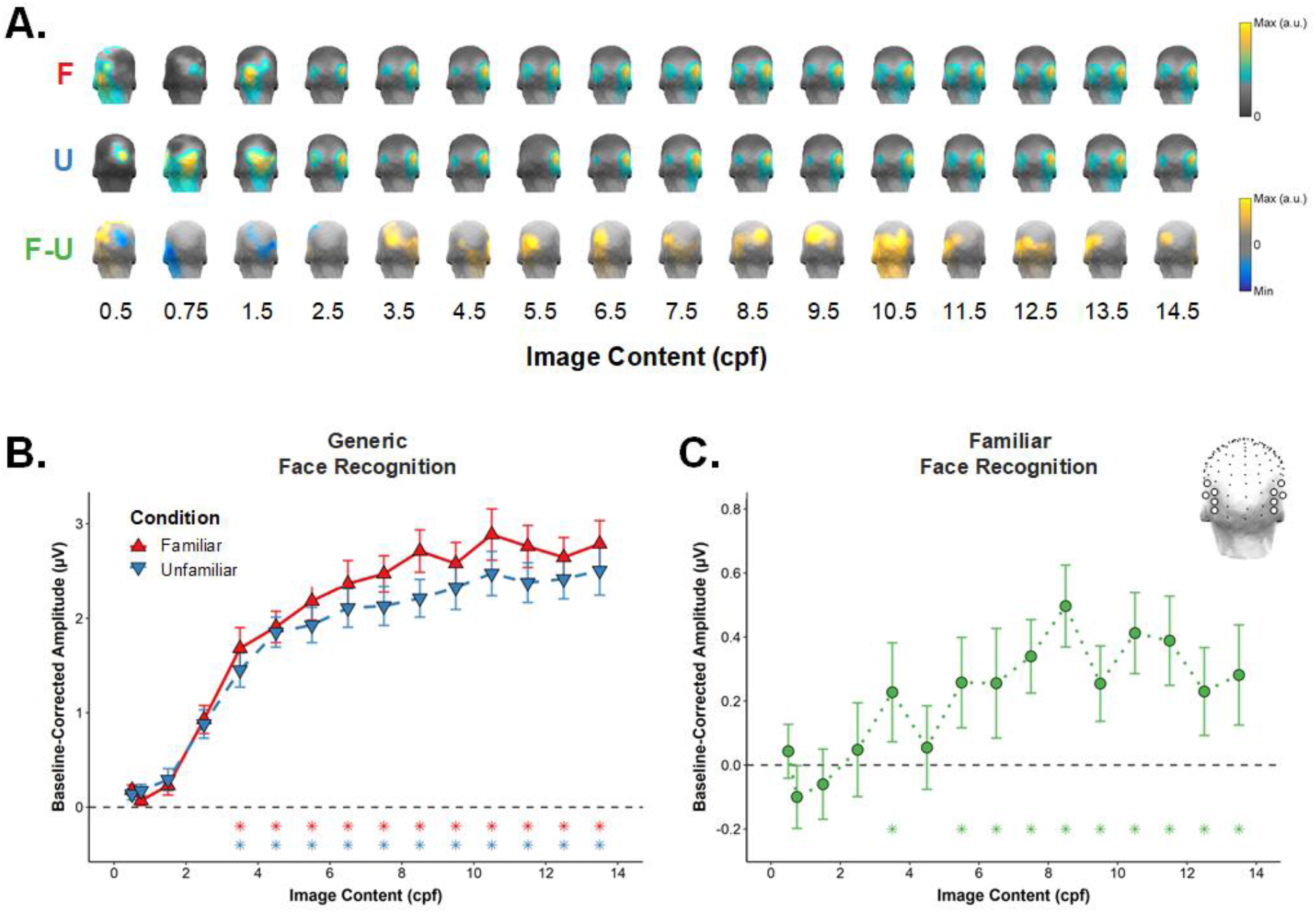
Group-level data for Expt. 2. **A.** Scalp topographies normalised to highlight the peak response locations across cpf values. **C.** Conditional mean face categorisation response profiles within the OT ROI (shown as right inset). **D.** The familiar face categorisation response profile within the same ROI. Error bars are SEM, asterisks indicate SF cutoffs at which a significant response was identified (*p* <. 01, one tailed).

Fig. 3 shows the individual threshold distributions for generic and familiar face recognition in Expt. 2. Here we observed very similar patterns to those found in Expt. 1: *Gen*-Thresh was very similarly distributed for the *Familiar* and *Unfamiliar* conditions, peaking in both cases over a narrow range of 2.5-4.5 cpf (5.3-9.5 cpi) (Wilcoxon *p*-value = 0.572; K-S *p*-value = .803). Interestingly, while the *Fam*-Thresh distribution was centred over the same cpf range as *Gen*-Thresh (Wilcoxon *p*-value = .109), its shape was noticeably different (K-S *p*-value = .036), being much wider and characterised by a long rightward tail. Overall, these results point to greater inter-observer variability in the image resolution needed to perceive the familiarity status of a face than to perceive its category. While coarse information may be sufficient for some observers, many require finer details to reliably recognise familiar faces.

We next evaluated whether there were general inter-individual differences affecting the efficiency of both generic and familiar face categorisation processes. As for Expt. 1, however, there was no evidence of a systematic relationship between individual participants’ *Gen*-Thresh and *Fam*-Thresh values (*r*pearson = 0.15, *p* = 0.52). As can be seen in Fig. 4B, participants for whom generic face categorisation was possible based on extremely coarse visual input did not also systematically recognise face familiarity at a comparatively coarse level. Finally, inspection of the topographical distribution at the individual observer categorisation thresholds revealed the similar profile as obtained in Expt. 1 (Fig. 5D). While statistical testing failed to reach significance threshold, *t*(1,17) = 1.56, *p* = 0.14; Fig. 5E), SF content levels enabling a distinction between *Familiar* and *Unfamiliar* faces also corresponded to a more bilateral response topography compared to SF steps at the generic face recognition threshold. This effect again appeared to be driven by more distributed hemispheric engagement when viewing *Familiar* faces compared to *Unfamiliar* ones (Fig. 5F).

## Discussion

Faces contain a wealth of information that appears available at once to the observer (e.g., sex, identity, emotion, etc.), yet little is presently known about how the brain achieves *concurrent* categorisation/recognition of these various face aspects. Here we provide the first systematic investigation of the relative informational dependencies underlying two concurrent and ecologically relevant recognition functions that follow a face encounter – generic face recognition and familiar face recognition. Using visual periodicity to isolate the selective neural response to unsegmented faces presented at regular intervals amid a wide variety of objects, we identified individual observer thresholds for each face function within two parametric manipulations of sensory input. In Expt. 1, manipulating image duration showed that exposures of just 33-50 ms enabled nearly all observers to consistently distinguish faces from a host of other categories (e.g., animals, plants, buildings, vehicles, etc.). In contrast, the temporal exposure observers required to recognise whether the faces were familiar was both higher on average and much more variable (*M* = 83 ms, range = 25-250 ms). In Expt. 2, manipulating image resolution yielded similar findings: nearly all participants recognised faces amongst objects based on extremely coarse visual input (i.e., just 2.5-4.5 cpf), but they exhibited considerable variability in how much finer resolution they required to recognise face familiarity (range = 1.5-13.5 cpf). In both experiments, the scalp distribution of neural responses was right-lateralised to a greater extent at the generic categorisation threshold than at the familiar face categorisation threshold. Critically, neither manipulation revealed a systematic offset in generic and familiar face categorisation across individual participants (i.e., observers who categorised faces vs. objects at short durations were not also able to recognise familiar faces at comparatively short durations). Together, these two lines of evidence indicate that during a face encounter, observers require less sensory evidence to successfully recognise that stimulus as a face than they do to recognise whether the face is familiar. In other words, generic face recognition is both more efficient, less variable, and processed in a more spatially constrained neural network than familiar face recognition.

Importantly, since the sensory input thresholds we report result from quantifying neural activation that generalises across a large number of highly variable, naturalistic exemplars (see examples in Figures 1 & S3), we can be confident that they relate to a form of face categorisation that goes beyond simple low-level featural differences between categories (25, 28). Moreover, in a crucial departure from previous efforts to compare the informational requirements of different types of face recognition across distinct task contexts and face presentations, the current thresholds pertain to categorisation functions elicited in parallel by *the same face encounters.* This notion of concurrence has gained recent traction in the face processing literature, with a number of recent studies using multivariate methods to probe the overlapping time-courses of various types of face categorisation reflected within the same neural response (i.e., decoding “which information is available when”, (12–15)). Here we took a very different approach, capturing differential responses to faces among objects regardless of exact timing (e.g., early/fast *vs.* late/slow). This enabled us to quantify how concurrent generic and familiar face recognition differed in their temporal/spatial sensory input requirements independently from any differences in their representational time-courses. In this way, our findings bring an important complementary perspective to the origin of putative processing speed differences between various dimensions of face recognition: Namely, if two components of face recognition differ in terms of how soon they can be achieved (7, 9) or decoded (14, 15) after face onset, this might not strictly reflect a difference in their relative speed of processing *per se*, but could equally arise if the processes unfold at the same speed, but with one requiring comparatively more evidence (i.e., signal-to-noise) than the other, and therefore being completed later (17). As such, our findings represent a vital step forward in understanding how face recognition operates in the real world, where in spite of vast differences in lighting, viewing angle, retinal size, and so on, a brief glance is all it takes to elicit rapid and simultaneous recognition of a face’s category, identity, sex, expression, and more.

### Face recognition based on brief image exposures

Although the present study is first to characterise how increasing image exposure duration affects *concurrent* forms of face categorisation/recognition, a number of studies taking a modular approach have considered the minimum viewing time these functions require separately (18–20). One such study that presented degraded synthetic faces at various brief durations found that observers required just 32ms image exposure to reach 75% accuracy for detecting whether a face was present/absent, but needed nearly double that time to reach the same threshold when identifying which of two facial identities had appeared (20). Our Expt. 1 findings accord well with these results in suggesting that sensory input demands are higher for more complex forms of face recognition. Remarkably, however, the same brief exposure durations that enable observers in this previous study to differentiate between faces and very simple distractors (i.e., scrambled face/oval outline) were also sufficient for our participants to recognise faces amid a rich variety of unsegmented objects (i.e., animals, plants, man-made artefacts, buildings, etc.). This highlights the efficiency of generic face recognition mechanisms, which appear capable of processing the highly complex visual information contained in natural images as well as simplistic features at very short image durations. In a similar vein, the critical image durations for generic face recognition we identify here neatly overlap those reported by a recent investigation of this face function that used frequency-tagging in the context of full-colour images and a completely orthogonal observer task (30) (i.e., ~33-50 ms *vs.* ~33-42 ms). Together with the reliable lower bound in individual observer thresholds in our Expt. 1, this consistency of image duration thresholds across differences in task and stimuli underscores the robustness and automaticity of generic face recognition in the human brain. Put differently, it appears there is indeed a hard limit to the minimum sensory input the human visual system requires to discern faces from other natural categories – one that does not seem lowered by attentional facilitation or richer image signal (31).

In contrast to the extremely brief exposures enabling successful generic face recognition, critical image durations previously implicated in identity-level face recognition are comparatively longer. For example, one study reported behavioural performance on a 6-way identity discrimination task to be at chance level for (masked) image durations of 17 ms and 33 ms, increasing markedly from durations of 50 ms upward (19). Elsewhere, the temporal processing capacity for face identification (one image per identity) has been estimated at 10 Hz, with one study demonstrating that 100 ms exposure provides sufficient inspection time for observers to distinguish recently learned faces from novel distractor ones (21). Yet since the duration threshold reported by this study corresponded to an arbitrary performance level (79%), we note that this estimate may in fact represent neither the minimal nor the optimal image duration for face identification. Crucially, to date no study manipulating viewing time has conclusively established the minimum sensory input requirements for successful recognition of *familiar* faces (i.e., famous faces). The data here indicate that this temporal threshold is on average very low (i.e., 50-83 ms), but also that the informational dependency of familiar face recognition is highly variable across individuals. While we can only speculate about the exact factors that drive this variability, presumably individuals vary in both the way they encode specific visual features of certain identities, and in the strength of more general semantic associations corresponding to that identity.

As a separate point, since the familiar face recognition threshold reported here results from quantifying neural activity selective for *either* familiar or unfamiliar faces presented amongst objects (and calculating a difference signal), it is important to note that this 50-83ms threshold range pertains to accessing face familiarity in the context of visually dissimilar object distractors. Distinguishing familiar and unfamiliar faces *directly* within a periodicity based design (e.g., UUUUFUUUUF…) would very likely demand a longer minimum exposure duration due the high stimulus similarity and resulting increased masking.

### Face recognition based on coarse visual input

The informational requirements for various face recognition functions have also been investigated by impoverishing face images by selectively removing content at certain spatial scales (i.e., spatial frequencies, SFs). Although the critical SF bandwidths considered diagnostic for different aspects of face recognition have received much attention (32, 33), to date these comparisons have largely focused on the finer-grained aspects of face perception (e.g., identity, emotional expression, gender, etc.). By comparison, investigations of critical SFs for recognising a face *as a face* have been much rarer (34). Our Expt. 2 data break new ground in this respect, by establishing the relative involvement of SF content for both the basic recognition and finer recognition of a face within the same individuals.

In terms of generic face recognition, the ultra-coarse information threshold in Expt. 2 (i.e., 2.5 – 4.5 cpf) is consistent with modular studies that have emphasised the importance of low SFs for this function (35, 36), albeit using more simplistic indices of this process (e.g., contrasting faces with just a single other category, such as cars). To date, only one other study has manipulated SF content for generic face recognition in the context of multiple naturalistic visual categories (22); previous work from our lab has demonstrated that this face function can be achieved based on images containing less than 5 cycles per image (corresponding to < 2 cpf). Together with the current results, these findings strongly suggest that discriminating faces from objects in natural environments depends on detecting the global structure of a face, rather than individual facial features which are not yet evident at the coarse spatial resolution of threshold images (see Fig. 4D).

The contribution of different spatial scales to familiar face recognition has received greater attention and there is general agreement that the critical SF bandwidth for identity-related face tasks comprises midrange values between 8-16 cpf (37). However, these studies have largely focused on identifying the *optimal* SF range for familiar face recognition, contrasting performance across different bandpass-filtered versions of the same face stimuli. By contrast, here we identified the *minimal* amount of cumulatively integrated SF content (across increasingly sharpened images) that enabled successful familiar face recognition. This approach revealed an unprecedented overlap in the coarse spatial resolutions (i.e., ~3.5 cpf) capable of driving both successful generic and familiar face recognition. Importantly, however, the fact that familiar face recognition *can* proceed based on highly degraded visual input does not imply that this process is optimally subserved by such limited image information. Rather, in the same way that finer-scale information improves the recognition of the stimulus as a face (38), the addition of high SF details likely serve to refine face identity representations, accentuating the difference between familiar and unfamiliar faces (39). Indeed, the rising recognition response profiles in our own data suggest this may indeed be the case.

### Broader implications & future directions

At a broader level, the current findings can be contextualised within the coarse-to-fine visual processing framework (33, 39–41), wherein limiting the temporal exposure of full-spectrum (i.e., unfiltered) images essentially serves to constrain the spatial scales the observer is able to extract while the image is onscreen. In this way, our two manipulations of face information and resulting findings potentially represent two sides of the same coin: upon encountering a face, the human visual system may rapidly extract its (coarse) global structure that suffices to distinguish it from other object categories, with more fine-grained sensory cues that facilitate familiarity recognition being more gradually accumulated. It is important to note, however, that the threshold offset between the two recognition functions should not imply that these phenomena represent different points on the same general face recognition evidence accumulation profile. Indeed, processes supporting the two functions may operate in parallel, and could potentially even be sub-served by different neural substrates, as their dissociation in cases of acquired prosopagnosia alludes to (42). Lateralisation differences in the current data hint at this possibility, insofar as distinguishing faces from objects mainly activated right-lateralised posterior sites, while recognising face familiarity recruited a more bilateral network. Future studies combining the current approach with intracerebral recordings or fMRI (28) will be important to clarify this issue.

From another perspective, the data here undermine the intuitive notion that in order to know whether a face is familiar or not, you must presumably already know that the stimulus before you is a face. This *basic-before-subordinate* notion (43) underlies many of the classic theoretical accounts of face recognition, which is claimed to unfold in a serial fashion from visual analysis (e.g., structural encoding of features) all the way up to the retrieval of high-level semantic associations (e.g., the name and context associated with that identity) (44, 45). By starting with the premise that a face is present in foveal vision, these models tacitly imply that low-level detection (e.g., figure-ground segmentation) and recognition of face category have already taken place. Yet our findings suggest that for a subset of observers, the same informational content can give rise to both successful generic and familiar face recognition of a face (46), suggesting that generic face recognition itself is a nontrivial stage that should be incorporated into face models.

Finally, although we might have expected to find a systematic relationship between an individual observer’s sensory input thresholds, observers in both experiments in fact exhibited highly consistent thresholds for generic face recognition compared with more variable thresholds for familiar face recognition. To the best of our knowledge, this relative difference in individual variance is a novel empirical observation with important implications for the neurofunctional basis of each process. That generic face recognition proceeds based on extremely impoverished information underlines the “hardwired” nature of this function – a process that both develops early (47) and appears robust to brain lesions that otherwise severely impair familiar face recognition (42). In contrast, the maturation of face identity processing, including familiar face recognition, is still much debated (48, 49) given the wide inter-individual variability in this function (50, 51). Related to this point is the fact that a small subset of our sample exhibited a lower sensory threshold for successful familiar face recognition than generic face recognition. Although such “negative” offsets may simply reflect signal-to-noise fluctuations associated with quantifying responses at the single-subject level, an interesting alternative might be that some individuals are able to detect the familiarity of a stimulus regardless of its visual category. Ascertaining whether these patterns represent a true functional phenomenon will necessitate probing the functional relevance of the neural thresholds reported here, ideally by relating them more directly to behavioural outcomes.

## Conclusion

Effective social behaviour in the real world depends on our ability to efficiently categorise faces along multiple dimensions at once (e.g., sex, emotion, identity, etc.), yet how the human brain achieves these concurrent categorisations is still not fully understood. Here we characterized the relative informational dependencies – quantified in space and time – of two critical and concurrent functions that follow a face encounter. Our data suggest that the sensory evidence the human brain requires to distinguish faces from other categories is systematically lower than that required to recognise faces we know. These findings underscore the neurofunctional distinctions between these two recognition functions - generic face recognition being robust in terms of its minimal sensory input requirements and low inter-individual variability, relative to the more demanding and idiosyncratic processes of familiar face recognition.

## Materials and Methods

### Participants

We tested independent samples of 25 participants each in Expts. 1 and 2. All gave written informed consent in accordance with UCLouvain BioEthics committee guidelines and were monetarily compensated for their time. All participants were right-handed, with normal or corrected-to-normal vision, and did not report any psychiatric or neurological history. All were of Belgian-French background or verified to be familiar with its culture. For each experiment separately, we excluded participants with poor behavioural performance (e.g., task performance < 2.5 *SD* from group average, details below) or those who blinked excessively during EEG recording (i.e., mean blinks / second > 2.5 *SD* from group average). The group-level accuracy and blink statistics were calculated on the full sample and both exclusion criteria were applied in parallel. The final sample consisted of 22 participants in Expt. 1 (13 females, mean age = 22.27 yrs ±1.98) and 21 participants in Expt. 2 (12 females, mean age = 21.81 yrs ±1.65).

### Protocol & Design

#### Expt. 1: Increasing Image Duration

Sequences contained face and object images whose presentation duration parametrically increased over the course of 84 seconds (Fig. 1B). Image duration began at 8.33 ms (i.e., 120 Hz = 120 images/s) and increased every 7 seconds across 12 steps until 333.33 ms (3 Hz). The SOA between face images was always 1000ms (i.e., face presentation frequency = 1 Hz), with the number of intervening objects between consecutive faces varying as a function of presentation duration step. The first and last sequence steps each lasted one extra second, during which the global image contrast gradually ramped up and down respectively. We included these “fade-in” and “fade-out” periods to reduce the impact of muscle artefacts and eye movements on the extremes of the EEG trace, and excluded these time-windows from response quantification. 24 sequences (12 *Familiar* and 12 *Unfamiliar*) were presented in pseudo-random order across two blocks whose order was counterbalanced across participants. At the start of the experiment, we ran an additional *Familiar* practice sequence to ensure participants understood task instructions^‡^.

#### Expt. 2: Increasing Image Content

Sequences contained face and object images with a fixed image duration of 83.33ms^§^ (i.e., 12 Hz = 12 images/s) whose spatial frequency (SF) content parametrically increased across 16 steps (Fig. 1C). Each step lasted 6 s, such that initially blurry images progressively sharpened over the course of 96 s (for movie of a similar SF sweep manipulation, see (22)). The initial step lasted for an extra 6 s, during which the global image contrast gradually ramped up in a “fade-in” period that was not included in analysis. In contrast to Expt. 1, here there were always exactly 8 object images between faces, for a fixed inter-face interval of 752 ms (i.e., face presentation frequency = 1.33 Hz). Observers saw 12 *Familiar* and 12 *Unfamiliar* sequences, presented in a pseudo-random order in two counterbalanced blocks.

#### Post-Sequence Identity Recognition Task

To ensure observers remained attentive during the image sequences and performed the categorisations of interest (i.e., focusing on faces among objects, and the identity of these faces), participants in both experiments completed a 2AFC identity recognition task after each sequence. This task comprised a 3 second display containing a probe face and foil face, whose positions were counterbalanced across sequences. Within the display duration, participants had to indicate with the left or right arrow key which of the two identities had appeared during the preceding sequence. The probe and foil faces always belonged to the same familiarity category as the preceding sequence (i.e., either both familiar or both unfamiliar), so participants could not respond based on a greater overall sense of familiarity. We emphasised to participants that they should respond based on the *identities* of the two individuals, not the specific images themselves (which were always novel). Task performance validated our familiarity manipulation, in that probe identification rates were significantly higher for familiar faces compared to unfamiliar ones in both Expt. 1 (F = 0.98±0.01; U = 0.79±0.03; *t*(21) = 6.87, *p* < 0.0001) and Expt. 2 (F = 0.98±0.60; U = 0.60±0.03, *t*(20) = 11.39, *p* < 0.0001). Response times were also faster for familiar compared to unfamiliar faces in Expt. 1 (F = 1.49±0.10; U = 2.01±0.10; *t*(21) = −7.05, *p* < 0.0001) and Expt. 2 (F = 1.52±0.11; U = 2.21 ±0.12, *t*(20) = −5.09, *p* < 0.0001).

### Stimuli & Display

Stimuli were 200 greyscale images of various non-face visual categories (e.g., animals, plants, structures, vehicles, objects, etc., 25, 27) and 240 greyscale images of faces (all unsegmented). Specific face images were selected on the basis of an independent stimulus pre-screening procedure (see Supp. Materials). *Familiar* identities were five international male actors/politicians considered highly recognisable to our Belgo-French population (George Clooney, Leonardo DiCaprio, Emmanuel Macron, Danny Boon, Nicolas Sarkozy). To ensure a similar overall level of photo quality, attractiveness, pose, etc., we matched each *Familiar* identity with an international celebrity of similar age/appearance whose recognition rate was close to chance for our population (respectively, Alfonso Cuarón, Thomas Kretschmann, Kirill Safonov, Najib Amhali, Humberto Zurita). For each of the resulting 10 identities, we sourced 24 exemplars that varied widely in pose, background, expression, lighting, viewpoint etc. The final 120 *Familiar* and 120 *Unfamiliar* faces were each divided into three subsets of 40 faces containing 8 exemplars per identity. During the two experiments, each of the three face subsets served as the face stimuli for four sequences (two in each testing block).

All object and face stimuli were sized 256×256 pixels and equalised in terms of mean luminance and contrast using the SHINE toolbox (52); this finalised image set comprised the stimuli used in Expt. 1. For Expt. 2 we took the additional step of generating spatially filtered versions of all images at 16 increasing low-pass filter cut-off values ranging from 1.05 to 30.6 cycles per image (cpi), corresponding to 0.5 to 14.5 cycles per face (cpf; estimated using the median width of faces within each image) (see Fig. 1C). We determined this range to encompass SF bands previously implicated in both generic face recognition (22) and face identity recognition (i.e., ~8-12 cpf) (37). The filter cut-off values were spaced to maximise resolution in cpf with the goal of identifying more precise thresholds at the individual subject level.

We used custom Java software to display stimuli on a 120Hz BenQ LED monitor with 1920×1080 resolution in a dimly lit room. The viewing distance was 50 cm, such that images spanned a visual angle of ~10°. All stimuli and instructions appeared on a grey uniform background; during the sequence presentation a small central fixation cross remained overlaid on the images.

#### Post-sequence 2AFC task

Stimuli for the 2AFC post-sequence identity recognition task were 24 unique probe + foil combinations (12 *Familiar* and 12 *Unfamiliar*). Each of the experimental 10 identities appeared as the probe at least twice, and was always paired with a novel, never-before-seen identity as the foil. Note that the actual images used for the experimental and 2AFC task were completely distinct, i.e., the probe was always a completely novel image of a previously-seen identity.

### EEG Acquisition & Analysis

We used a BioSemi Active 2 system (Amsterdam, Netherlands) with standard 10–20 system electrode locations and additional intermediate positions to acquire 128-channel scalp EEG. We monitored eye movements using electrodes placed at the outer canthi of both eyes, as well as above and below the right eye. Data was sampled at 512 Hz and individual electrode offsets were held below ±50 μV. During testing, digital triggers were sent via a parallel port at the start and the end of each stimulation sequence, and when the participant made a behavioural response. The experimenter manually initiated the recording for each sequence only after the EEG trace showed no muscular/ocular artefact for at least 5 s. Participants took a short break after every six sequences to rest their eyes.

#### EEG Preprocessing

We analysed EEG data offline using Letswave 5 (https://www.letswave.org/) running on MATLAB R2012b (MathWorks, MA, United States). We realigned the continuous EEG data to remove abrupt signal offsets that resulted from pausing the recording, then de-trended and removed the DC component from the data. Next we applied a band-pass filter with cut-offs at 0.05 Hz and 125 Hz (4^th^ order zero-phase Butterworth filter), followed by a multi-notch filter remove electrical noise carried at 50, 100, and 150 Hz (FFT filter, width = 0.5). Data was downsampled to 256 Hz for easier handling and storage, and segmented according to stimulation sequences, with two extra seconds before and after the sequence (Expt. 1 = 86 s; Expt. 2 = 102 s). For each participant, we used independent component analysis (ICA) with a square mixing matrix to remove a single component corresponding eyeblinks (identified through visual inspection of component waveforms and topographical distributions). We interpolated artefact-ridden channels with the average of the 3 neighbouring channels (less than 5% of channels were corrected for each observer) and re-referenced the cleaned data to the average of all 128 scalp channels. We cropped the preprocessed data to exclude the fade-in and fade-out periods for each sequence (final epoch lengths were 84 s in Expt. 1; 96 s in Expt. 2). We then averaged each participant’s *Familiar* and *Unfamiliar* segments separately, before chunking these conditional averages into separate epochs for each duration/SF step (Expt. 1 = 12 x 7 s epochs, Expt. 2 = 16 x 6 s epochs). Finally, we applied a Fast Fourier Transformation (FFT) to each epoch to extract 0-128 Hz frequency amplitude spectra for each combination of participant, condition, and step (frequency resolution in Expt. 1 = 0.14 Hz, in Expt. 2 = 0.16 Hz). We considered conditional group means within a predefined bilateral OT region-of-interest (ROI), averaging across electrode sites previously shown to be involved in both face categorisation and recognition (P8, PO8, P10, PO10, PO12, P7, PO7, P9, PO9, PO11, see insets on Figures 2 & 6). All group-level and individual response profiles are noise-corrected amplitudes, obtained by subtracting the mean noise value from the signal, separately for each condition and each step.

#### Threshold Analysis

To obtain face recognition thresholds at both the group and individual level, we used a bootstrapping procedure in which we tested the magnitude of the relevant signal against an empirical EEG noise distribution. To determine the generic face recognition threshold, we first obtained a signal estimate by summing the amplitude values at the harmonics of the face presentation frequency up to 30 Hz (Expt. 1: 30 harmonics = 1-30 Hz; Expt. 2: 22 harmonics = 1.33-29.26 Hz), separately for each step/condition combination. Harmonics overlapping those of the image presentation frequency were not included in this summation process. Next, we computed a noise estimate by summing the same number of amplitude values at randomly selected frequencies, excluding the image and face presentation harmonics. In order to account for the 1/f profile of the EEG spectrum, the frequency range for the noise calculation was slightly expanded (3 extra frequency bins above and below the face categorisation response range; Expt. 1 = 0.58-30.42 Hz; Expt. 2 = 0.85-29.74 Hz). We repeated this noise estimation procedure 10,000 times to generate a noise distribution. At each combination of condition/step, we considered the generic face categorisation response to be significant if it exceeded the top 1% of its corresponding noise distribution (i.e., *p* < 0.01), defining the threshold for this function (*Gen*-Thresh) as the first step at which this significance criterion was reached. To determine each observer’s familiar face recognition threshold (*Fam-*Thresh), we applied the same procedure to the difference amplitude spectrum of the two conditions (i.e., Familiar– Unfamiliar) and used the same 99^th^ percentile cut-off. The direction of this subtraction pinpoints the step at which the face-selective response elicited by familiar faces was larger than that elicited by unfamiliar faces.

We compared the resulting threshold distributions using two nonparametric statistical tests: the Wilcoxon matched-pairs signed rank test, which examines the central tendency across distributions (i.e. null hypothesis supposes the same mean/median values), and the two-sample Kolmogorov-Smirnov (K-S) test, which considers the equality of empirical cumulative distribution functions (i.e., takes into account the global shape of the distributions). To circumvent the problem of ties in the data (differences of zero that preclude the calculation of exact *p*-values), we added a very small amount of jitter to each value in the test vectors (+/- < 0.01 ms or cpf), and repeated this process 10,000 times. In both the Wilcoxon and the K-S tests, differences were deemed significant if the mean *p-*value obtained across these 10,000 iterations was <0.05; in the results we report these mean *p*-values.

#### Lateralisation analysis

Hemispheric differences at thresholds were examined only for observers for whom both *Gen-* Thresh and *Fam-*Thresh could be defined (Expt. 1: N = 20; Expt. 2: N = 18). For each observer, we isolated the duration/SF content step corresponding to their individually-defined threshold for generic or familiar face recognition. We then averaged across the data in these individually-defined steps in two ways. To inspect the scalp topographies at *Gen*-Thresh, we averaged each participant’s *Familiar* and *Unfamiliar* data and calculated a group mean from the resulting averages. To inspect the scalp topographies at *Fam*-Thresh, we performed the *Familiar – Unfamiliar* subtraction for each participant and averaged across the resulting differences. Next we quantified the degree of lateralisation at each threshold point by calculating the difference between corresponding right and left electrode sites within our *a priori* OT ROI (see above), comparing these lateralisation indices at *Gen*-Thresh and *Fam*-Thresh using a two-tailed pairwise *t*-test. Lastly, to examine how potential lateralisation differences arose, we further decomposed the responses at each threshold by averaging neural activity separately for *Familiar* and *Unfamiliar* conditions.

## Acknowledgments

We thank A. Conte for developing the Java stimulation software and V.Goffaux for providing the Matlab scripts used to generate the low-pass filtered images in Expt. 2.

## Footnotes

* The term “recognition” refers here to the production of a selective (i.e., discriminant) response to a given sensory input, a response that can be reproduced (i.e., generalized) across variable viewing conditions. In this sense, recognition is essentially a categorization function, and the two terms are used interchangeably in the manuscript.

† Our focus on *informational* dependencies should not imply that face categorization is governed solely by physical characteristics of visual input. Indeed, to any naïve system (human or artificial), there is no stimulus-level information that distinguishes a ‘familiar’ face from an ‘unfamiliar’ one. Here we assume that the integration of sensory information with other semantic/memory processes to be inherent to generic and familiar face recognition.

‡ Prior to the practice trial, both experiments contained an additional four sequences in which there was no parametric variation (i.e., observers saw full spectrum images presented at 12 Hz with a fixed face frequency of 1.5 Hz). These additional sequences pertained to a separate experimental investigation and are not reported further here.

§ We a fixed image presentation rate of 12 Hz in Expt. 2 since this stimulation rate has previously been shown to elicit clear face categorisation responses within individual observers (27)

