## Supplemental Materials for "Critical information thresholds underlying concurrent face recognition functions"

*Supplemental Fig. 1*

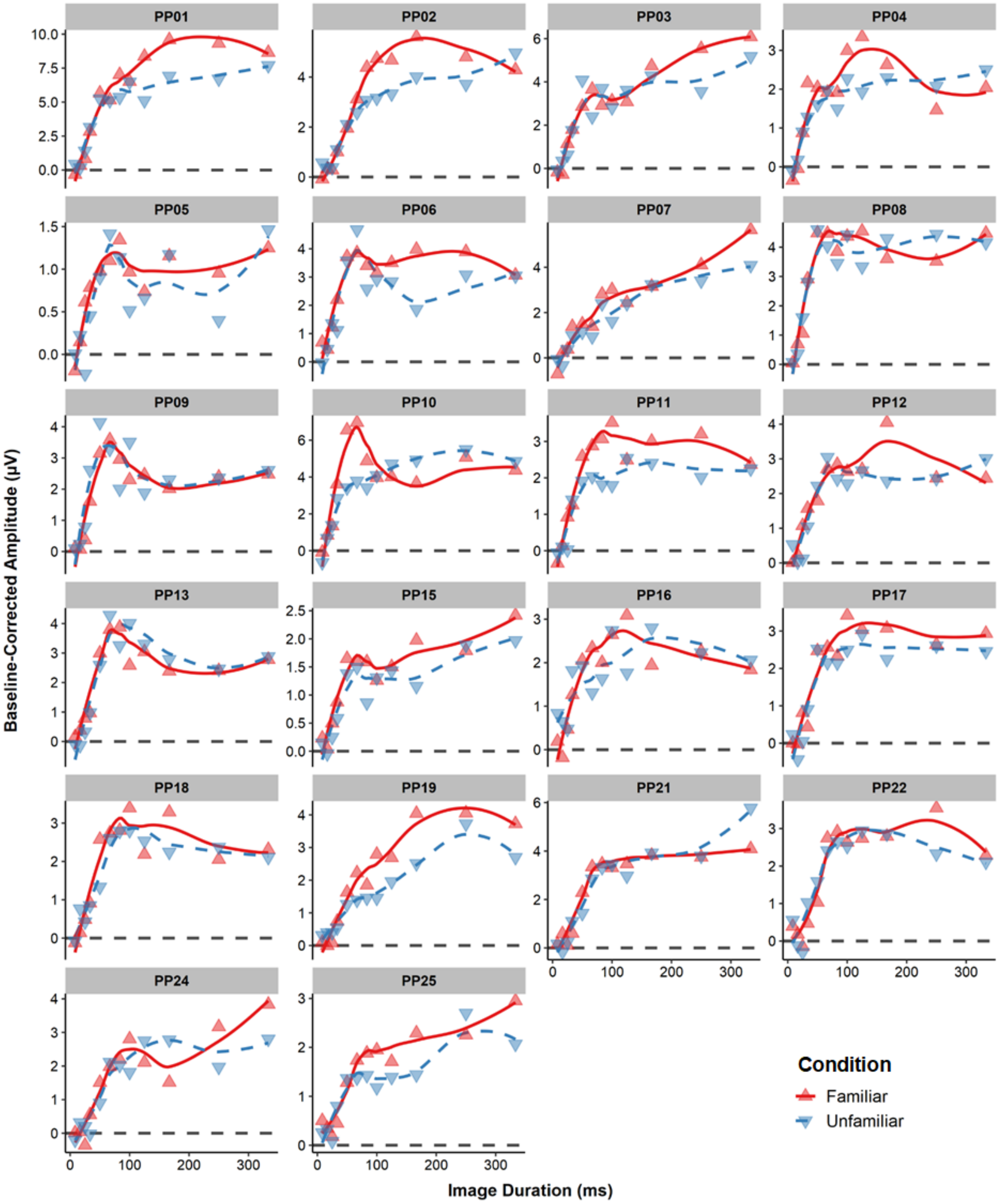


**Fig. S1**. *Familiar* and *Unfamiliar* face categorisation response profiles for individual participants in Expt. 1, shown with local polynomial regression fits. At the individual observer level, differences were generally transient rather than sustained as in the group-level analysis. Interestingly, the later steps corresponding to the longest image durations consistently showed the least differentiation between response functions for *Familiar* vs. *Unfamiliar* faces.

***Supplemental Fig. 2***

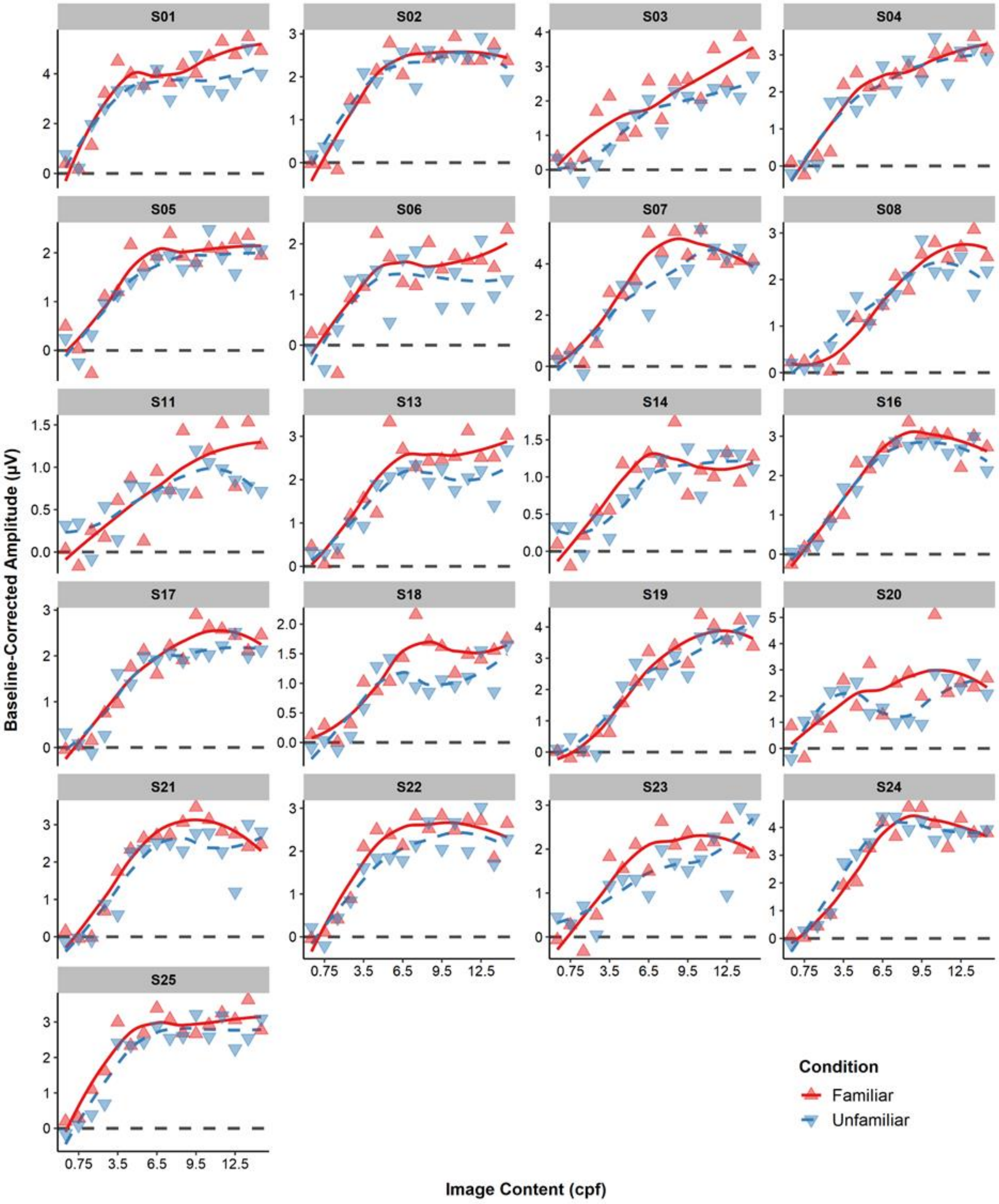


**Fig. S2**. *Familiar* and *Unfamiliar* face categorisation response profiles for individual participants in Expt. 2, shown with local polynomial regression fits. As for Expt. 1, differences between these profiles tended to be transient at the individual participant level, such that later steps containing the highest information content also tended to show the least differentiation between the *Familiar* and *Unfamiliar* profiles.

***Stimulus Pre-Selection Procedure***

Specific face identities and exemplar images were selected on the basis of an independent preselection procedure. We began by nominating 13 celebrities considered highly recognisable to a Belgo-French population. Each of these *Familiar* identities was paired with a comparatively unknown foreign celebrity of similar age and appearance. For each of these 26 identities, we sourced 30 individual exemplar images online (i.e., 780 images, each 256×256 pixels), ensuring the sets varied widely in terms of background, lighting, facial expression, age, pose, etc. (Fig. S3B). We used a browser-based experimental platform (https://www.testable.org), to present 50 Belgo-French participants aged between 18-42 years (18 males, mean age = 23 years±3.62) with a 2AFC naming task (380 trials) containing these images (i.e., each participant saw half the exemplar images for each identity). Each trial contained a brief fixation cross, followed by a central target face image (e.g., image of George Clooney) with a target and lure name below (e.g., “GEORGE CLOONEY” --- “BRAD PITT”, target name position randomised across trials). The face image disappeared after 500ms, while the names remained onscreen. Participants had a maximum of 3 seconds from display onset to click on the name that matched the face image on that trial. Both names were always drawn from the same Familiarity category (e.g., a famous face always appeared with two famous names), to prevent participants from responding based on a sense of familiarity alone.

The five identities with the highest recognition rates were Danny Boon, Nicolas Sarkozy, George Clooney, Leonardo DiCaprio, and Emmanuel Macron ($M$ = 96.55%, $SD$ = 0.91). Recognition rates for their corresponding matched identities were much lower and close to chance ($M$ = 60.51%, SD = 3.47), $t(4) = 27.68$, $p < .0001$, and there was a significant difference in response time between the *Familiar* (M = 1216 ms, SDF = 30 ms) and *Unfamiliar* identities ($M$ = 1488 ms, SD = 62 ms), $t(4) = -15.88$, $p < .0001$. Since the familiar identities in this group were both *i)* highly recognisable, and *ii)* very distinct from their unfamiliar counterparts, we selected these 10 face identities to use in the main experiments, and kept the remaining identities aside to use as foil faces during the post-sequence 2AFC identity recognition task.
